# CUL-6/cullin ubiquitin ligase-mediated degradation of HSP-90 by intestinal lysosomes promotes thermotolerance

**DOI:** 10.1101/2023.12.21.572895

**Authors:** Mario Bardan Sarmiento, Spencer S. Gang, Patricija van Oosten-Hawle, Emily R. Troemel

**Author notes:** Correspondence may be addressed to Emily Troemel.

## Abstract

Heat shock can be a lethal stressor. Previously, we described a CUL-6/cullin-ring ubiquitin ligase complex in the nematode *Caenorhabditis elegans* that is induced by intracellular intestinal infection and proteotoxic stress, and that promotes improved survival upon heat shock (thermotolerance). Here, we show that CUL-6 promotes thermotolerance by targeting the heat shock protein HSP-90 for degradation. We show that CUL-6-mediated lowering of HSP-90 protein levels, specifically in the intestine, provides a thermotolerance benefit independent of heat shock factor HSF-1. Furthermore, we show that lysosomal function is required for CUL-6-mediated promotion of thermotolerance and that CUL-6 directs HSP-90 to lysosome-related organelles upon heat shock. Altogether, these results indicate that a CUL-6 ubiquitin ligase promotes organismal survival upon heat shock by promoting HSP-90 degradation in intestinal lysosomes. Thus, HSP-90, a protein commonly associated with protection against heat shock and promoting degradation of other proteins, is actually itself degraded to protect against heat shock.

## Introduction

Exposure to dangerously high temperatures, i.e. heat shock, can lead to organismal sickness and death. These negative impacts are due, at least in part, to heat-induced denaturation and aggregation of proteins, which impair normal cellular function. To combat these negative impacts, organisms have evolved dedicated stress resistance pathways such as the heat shock response, which upregulates and deploys heat shock proteins (chaperones) to refold denatured proteins and restore protein homeostasis (proteostasis) in response to acute increases in temperature.^1,3^ A central regulator of the heat shock response and proteostasis is the transcription factor Heat Shock Factor 1 (HSF-1), which mediates the transcriptional upregulation of chaperones upon heat shock to promote thermotolerance.^4-6^ HSF-1 is conserved across organisms from yeast to the nematode *C. elegans* to humans.

In C. *elegans*, we recently described the Intracellular Pathogen Response (IPR) as a stress resistance pathway that appears to promote thermotolerance in a manner that is separate from the upregulation of chaperones by HSF-1.^7-10^ The IPR comprises a common set of genes induced by natural intracellular pathogens of the intestine, including microsporidia and the Orsay virus, as well as by abiotic stressors like proteasome blockade and chronic heat stress.^11^ A negative regulator of the IPR is *pals-22*, a protein of unknown biochemical function but nonetheless serves a critical physiological function to repress IPR mRNA expression in the absence of infection or stress. Constitutively upregulated IPR gene expression in *pals-22* mutants causes slowed development but increased resistance to infection and heat shock.^7 11 12^

IPR genes are not enriched for chaperones and instead are enriched for transcriptionally upregulated E3 cullin-ring ubiquitin ligase components, which we found are required for the increased thermotolerance of *pals-22* mutants.^8,11^ E3 ubiquitin ligases are enzymes that conjugate the small protein ubiquitin onto lysine residues of substrate proteins, which then alters the fate of these proteins.^13^ Skp-Cullin-F-box (SCF) ubiquitin ligases are a large class of cullin-ring ubiquitin ligases and are multi-subunit enzymes composed of three core components: 1) Cullins, 2) Skp proteins and 3) RING proteins.^14^ These core components assemble with F-box proteins that serve as adaptors to recognize substrate proteins for ubiquitylation. IPR-induced SCF genes include cul-6/cullin, as well as a previously uncharacterized RING protein *(rcs-1)*, three Skp-related proteins *(skr-3, skr-4, skr-5)* and two previously uncharacterized F-box proteins *(fbxa-75, fbxa-158)*. Using genetics and biochemistry, we demonstrated that CUL-6 protein assembles with these other SCF protein components into a multi-subunit ubiquitin ligase complex that promotes thermotolerance in *pals-22* mutants.^15^ Both *pals-22* and *cul-6* are normally expressed in the intestine, among other tissues, and expression of *pals-22* or *cul-6* only in the intestine can regulate thermotolerance.^7,15^

The ubiquitin ligase activity of cullins can be increased by post-translational modification of a conserved lysine by the ubiquitin-like protein NEDD8 in a process called neddylation.^16^ We found that the ability of CUL-6 to promote thermotolerance in *C. elegans* depends on a conserved neddylation site,^15^ indicating that the CUL-6 ubiquitin ligase complex likely promotes thermotolerance through its ability to conjugate ubiquitin onto substrates, but it was unclear what these substrates were. One hypothesis was that a CUL-6 ubiquitin ligase could target misfolded proteins for destruction, including pathogen proteins delivered into host cells in the context of infection, and/or that it might target misfolded cytosolic host proteins in the context of heat shock.^8^ The ultimate fate of CUL-6 target proteins was also not clear. The most common fate for ubiquitylated proteins is degradation by the proteasome, but the lysosome can also degrade them,^17^ or ubiquitylation can result in non-degradative effects such as altered trafficking, subcellular localization, or biochemical function.^18^

Here, we provide evidence that the highly abundant heat shock protein HSP-90 is a target for degradation by the CUL-6 ubiquitin ligase complex. Unlike other heat shock proteins, HSP-90 is highly expressed in the absence of heat shock and has many functions, including facilitating the proper folding of hundreds of proteins under unstressed conditions.^19-21^ We find that CUL-6 activity reduces HSP-90 protein levels in the absence of heat shock, and that decreased expression of HSP-90, specifically in the intestine, leads to higher thermotolerance. In contrast, overexpression of HSP-90 in the intestine leads to lower thermotolerance. The current paradigm in the field indicates that loss of HSP-90 activates HSF-1,^22,23^ but here, we find that CUL-6-mediated effects of lowering HSP-90 levels on thermotolerance appear independent of HSF-1. To investigate where HSP-90 is degraded in the cell, we show that lysosomes regulate the levels of HSP-90 protein, and we show that the effects of HSP-90 and CUL-6 on thermotolerance depend on lysosomal function. Furthermore, we show that CUL-6 directs HSP-90 to lysosome-related organelles in the intestine upon heat shock. Altogether, our results support a model that the CUL-6 ubiquitin ligase targets HSP-90 for ubiquitylation, which leads to its subsequent degradation by the lysosome and/or lysosome-related organelles to promote organismal survival upon heat shock in an HSF-1-independent manner.

## Results

### The lysosome is specifically required for CUL-6-mediated thermotolerance as part of the IPR

Ubiquitylated substrates can be degraded by the proteasome or by the lysosome. To determine whether the proteasome or the lysosome might degrade the substrate(s) of the IPR-induced CUL-6 ubiquitin ligase, we blocked each pathway with pharmacological inhibitors while performing heat shock to assess changes in CUL-6-dependent thermotolerance. First, we treated wild-type animals or *pals-22* mutants (which have *cul-6* upregulated as part of the IPR and have increased thermotolerance) with the proteasome inhibitor bortezomib. Here, we found that bortezomib treatment impaired thermotolerance in both a wild-type and a *pals-22* mutant background, indicating proteasome blockade negatively affects heat shock survival in a CUL-6-independent manner (Figure 1A, S1A). In contrast, we found that treatment with the lysosome inhibitor bafilomycin impaired thermotolerance only in a *pals-22* mutant background and not in a wild-type background, suggesting that the lysosome is specifically important to promote CUL-6-mediated thermotolerance when the IPR is induced (Figure 1B).

**Figure 1.**
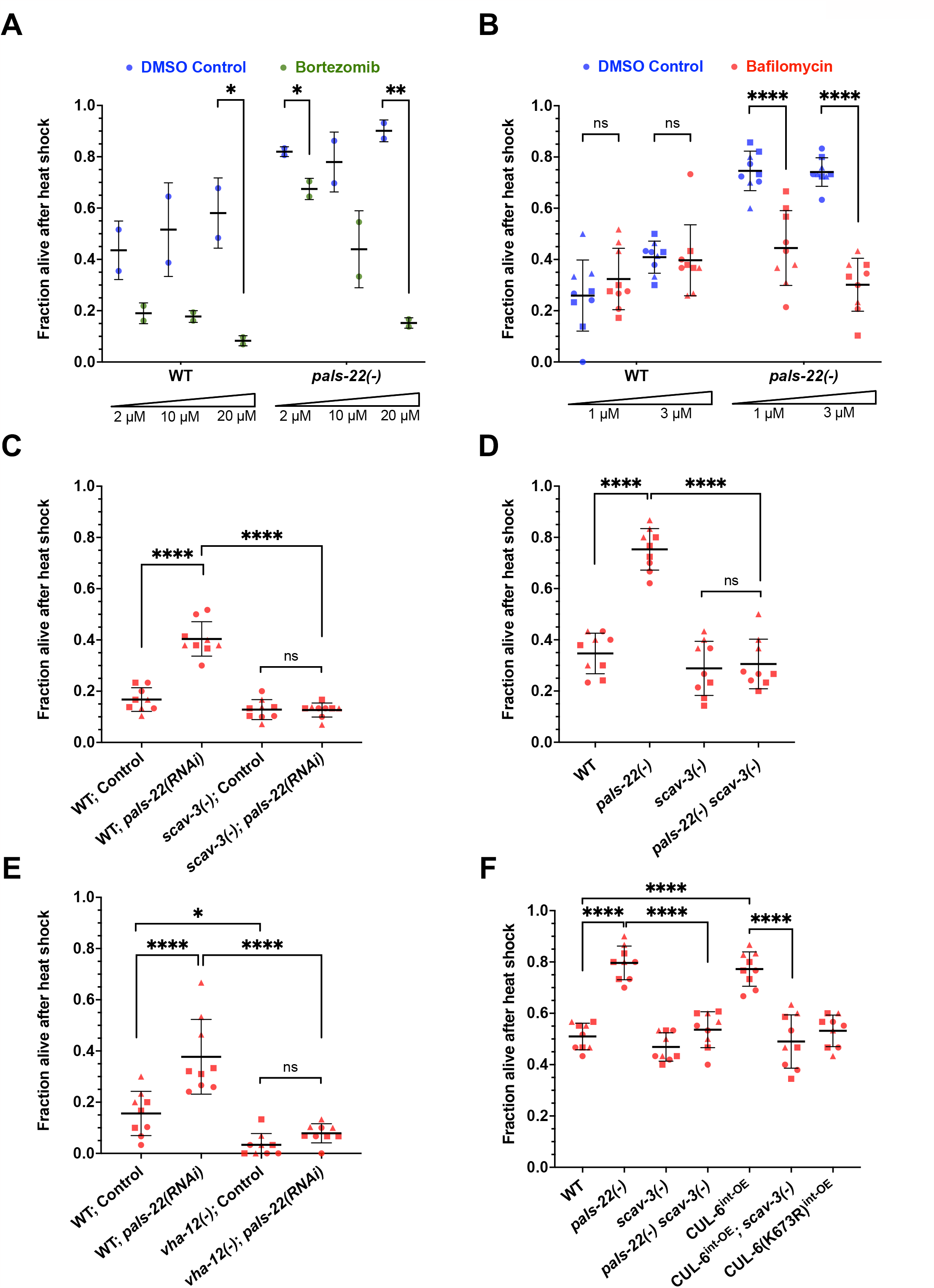
The lysosome is required for CUL-6-mediated thermotolerance as part of the IPR. (A, B) Survival of wild-type or *pals-22(-)* mutant animals treated with bortezomib (A) or bafilomycin (B) after 2 h of 37.5 °C heat shock with a 15 min gradual ramp-up, followed by a 24 h recovery period at 20 C (hereafter referred to as “heat shock treatment”). Bortezomib-treated animals (green dots) were tested with two plates over one experiment with 30 animals per plate, and bafilomycin-treated animals (red dots) were tested in triplicate experiments with three plates per experiment and 30 animals per plate. DMSO-treated animals (blue dots) were tested as the vehicle control for both inhibitors. Unpaired t-tests for each genotype and concentration of inhibitor were used to calculate p-values.(C) Heat shock treatment of wild-type or *scav-3(-)* mutant animals fed *E. coli* OP50 expressing control or *pals-22* double-stranded RNA (dsRNA) to induce RNAi (see methods). (D) Survival of wild-type, *pals-22(-), scav-3(-)*, or *pals-22(-) scav-3(-)* mutant animals after heat shock treatment. (E) Heat shock treatment of wild-type or *vha-12(-)* mutant animals fed OP50 expressing control or *pals-22* dsRNA to induce RNAi. (F) Heat shock treatment of animals from D, also including strains with transgenes over-expressing CUL-6 in the intestine (superscript int-OE). CUL-6(K673R) indicates a lysine to arginine mutation at CUL-6’s neddylation site with reduced ubiquitylation activity, which controls for CUL-6 overexpression in the intestine. For C-F, animals were tested in triplicate experiments with three plates per experiment and 30 animals per plate. A one-way ANOVA with Tukey’s multiple comparisons test was used to determine p-values. For A-F, the mean fraction of animals alive for the pooled replicates is indicated by the black bar with error bars as the standard deviation (SD). Each dot represents a plate, and different shapes represent the experimental replicates performed on different days. *p< 0.05, **p < 0.01, ****p < 0.0001.

Next, we tested the role of the lysosome in promoting thermotolerance using genetic approaches. Here, we performed RNAi against *pals-22* in animals with a mutation in *scav-3*, a lysosomal membrane protein important for lysosomal integrity.^24^ We found that *pals-22* RNAi, which increases thermotolerance in wild-type animals, no longer increased thermotolerance in *scav-3* mutants (Figure 1C). Similarly, *pals-22 scav-3* double mutants have a thermotolerance phenotype similar to that of wild-type animals (Figure 1D). Furthermore, *pals-22* **RNAi** no longer increased thermotolerance in mutants defective in *vha-12*, which encodes a Vacuolar-ATPase subunit important for lysosomal function^25^ (Figure 1E), again indicating that the lysosome is required for the increased thermotolerance in pals-22-defective animals.

The increased thermotolerance of *pals-22* mutants depends on *cul-6* expression in the intestine, and overexpression of *cul-6* specifically in the intestine, even in a wild-type background, promotes thermotolerance.^15^ Investigating the role of the lysosome in this context, we saw that the increased thermotolerance due to *cul-6* overexpression depended on *scav-3* (Figure 1F). As a control, we show that overexpression of a neddylation-deficient CUL-6(K673R) that lacks the lysine used for neddylation does not promote thermotolerance (Figure 1F), consistent with prior results.^15^ Taken together, these findings indicate that the lysosome is required for the increased thermotolerance associated with transcriptional upregulation of *cul-6*.

### RNAi knock-down of *hsp-90* specifically in the intestine increases thermotolerance independently of HSF-1

Given the results above indicating that lysosomal degradation of a CUL-6 ubiquitin ligase substrate promotes thermotolerance, we searched for candidate proteins whose reduction might promote thermotolerance. Previously published studies have shown that, in some cases, levels of the heat shock protein HSP-90 paradoxically appear to be negatively associated with thermotolerance.^23,26,27^ Therefore, we investigated the possible role of HSP-90 in our heat shock paradigm and indeed found that feeding animals *hsp-90* dsRNA significantly increased their thermotolerance (Figure 2A).

**Figure 2.**
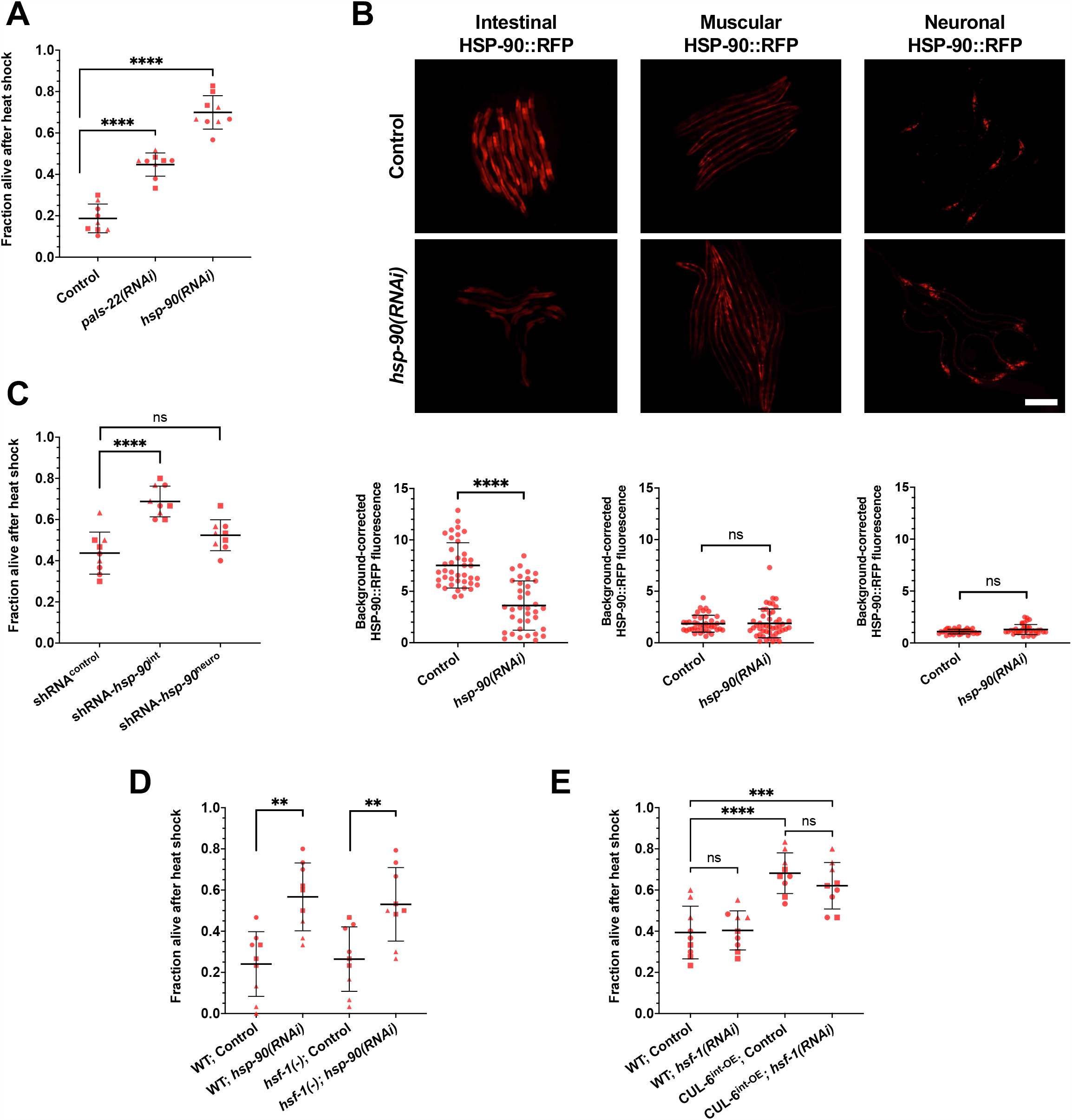
Knockdown of *hsp-90* in the intestine increases thermotolerance. (A) Heat shock treatment of wild-type animals fed OP50 expressing control, *pals-22*, or *hsp-90* dsRNA to induce RNAi. (B) Top; Representative fluorescent images of L4 animals with HSP-90::RFP overexpression under different tissue-specific promoters fed OP50 expressing control or *hsp-90* dsRNA to induce RNAi. Scale bar= 200 µm. Bottom; quantification of the RFP fluorescent signal for each condition. Each dot represents one animal measured, and the data shown are the results of three experimental replicates. n = 36 – 47 worms quantified. A Mann-Whitney test was used to calculate p-values for all strains. (C) Heat shock treatment of animals expressing tissue specific short hairpin RNA (shRNA) against *hsp-90* in the intestine and neurons relative to a control strain. (D) Heat shock treatment of wild-type or *hsf-1(-)* mutant animals fed OP50 expressing control or *hsp-90* dsRNA to induce RNAi. (E) Heat shock treatment of wild-type and CUL-6^int-OE^ animals fed OP50 expressing control or *hsf-1* dsRNA to induce RNAi. For A, C, D, and E, animals were tested in triplicate experiments with three plates per experiment and 30 animals per plate. The mean fraction of animals alive for the pooled replicates is indicated by the black bar with error bars as the SD. Each dot represents a plate, and different shapes represent the experimental replicates performed on different days. A one-way ANOVA with Tukey’s multiple comparisons test was used to calculate p-values. For A-E, **p < 0.01, ***p < 0.001, ****p < 0.0001.

To examine the efficacy of *hsp-90* RNAi in various tissues, we next performed whole-animal *hsp-90* RNAi by feeding dsRNA in transgenic strains that express HSP-90::RFP specifically in the intestine, the body wall muscle, or in neurons.^26^ Here, we only observed a significant reduction of HSP-90::RFP levels in the intestine (Figure 2B). In *C. elegans*, the neurons are often refractory to RNAi delivered by feeding, so the lack of knock-down there is not surprising.^28^ However, the muscle is generally susceptible to feeding **RNAi**. One potential explanation for this lack of effect is the compensatory mechanisms that have been reported to upregulate *hsp-90* expression after *hsp-90* RNAi in *C. elegans*,^26^ or after deleting 3 of the 4 *hsp-90* alleles found in mice.^29^ These compensatory mechanisms may be responsible for maintaining stable HSP-90::RFP protein levels in muscle after whole-animal RNAi in *C. elegans* (Figure 2B).

To knock-down HSP-90 in specific tissues, and to circumvent the issues with neurons being refractory to feeding RNAi, we next used strains that genetically encode short hairpin RNA (shRNA) against *hsp-90* specifically expressed in different tissues.^27^ Here, we found that by targeting *hsp-90* for RNAi specifically in the intestine there was increased thermotolerance (Figure 2C). In contrast, a strain with *hsp-90* shRNA expressed specifically in neurons did not exhibit significantly increased thermotolerance (Figure 2C). Altogether, these results demonstrate that reducing HSP-90 levels specifically in the intestine leads to increased thermotolerance.

Prior studies have indicated that HSP-90 binds to the transcription factor HSF-1 and holds it in an inactive state until heat shock, at which point HSP-90 binds to misfolded proteins and releases HSF-1 to activate the transcription of chaperones.^22^ Therefore, we examined whether HSF-1 was required for the increased thermotolerance either through the reduction of HSP-90 levels or the overexpression of CUL-6. First, we performed *hsp-90* RNAi in an *hsf-1* mutant background and saw increased thermotolerance similar to *hsp-90* RNAi in a wild-type background (Figure 2D). Of note, while the overexpression of *hsf-1* in *C. elegans* consistently promotes thermotolerance, there are varying results in the loss of *hsf-1* function; some studies show reduced thermotolerance and others show no reduction in thermotolerance, similar to our results here.^30-33^ Next, we performed *hsf-1* RNAi in a *cul-6* intestinal overexpression strain and saw no significant impairment on thermotolerance (Figure 2E). Therefore, the increased thermotolerance due to the effects of CUL-6 overexpression and loss of HSP-90 appear to be independent of HSF-1 and the canonical heat shock response.

### Overexpression of HSP-90 in the intestine decreases thermotolerance independently of HSF-1

Next, we investigated whether elevating HSP-90 levels in the intestine would impair thermotolerance. We performed heat shock assays on transgenic strains where HSP-90::RFP is overexpressed specifically in the intestine, body wall muscle, or neurons. Here we found HSP-90::RFP overexpression in the intestine decreased thermotolerance, while overexpression in body wall muscle and neurons did not negatively affect thermotolerance (Figure 3A). These results are consistent with a prior study that found HSP-90 overexpression in the intestine impaired thermotolerance.^26^ However, that study found overexpression of HSP-90 in muscle and neurons also impaired thermotolerance, which may be due to distinct strains and assay conditions used in that study.

**Figure 3.**
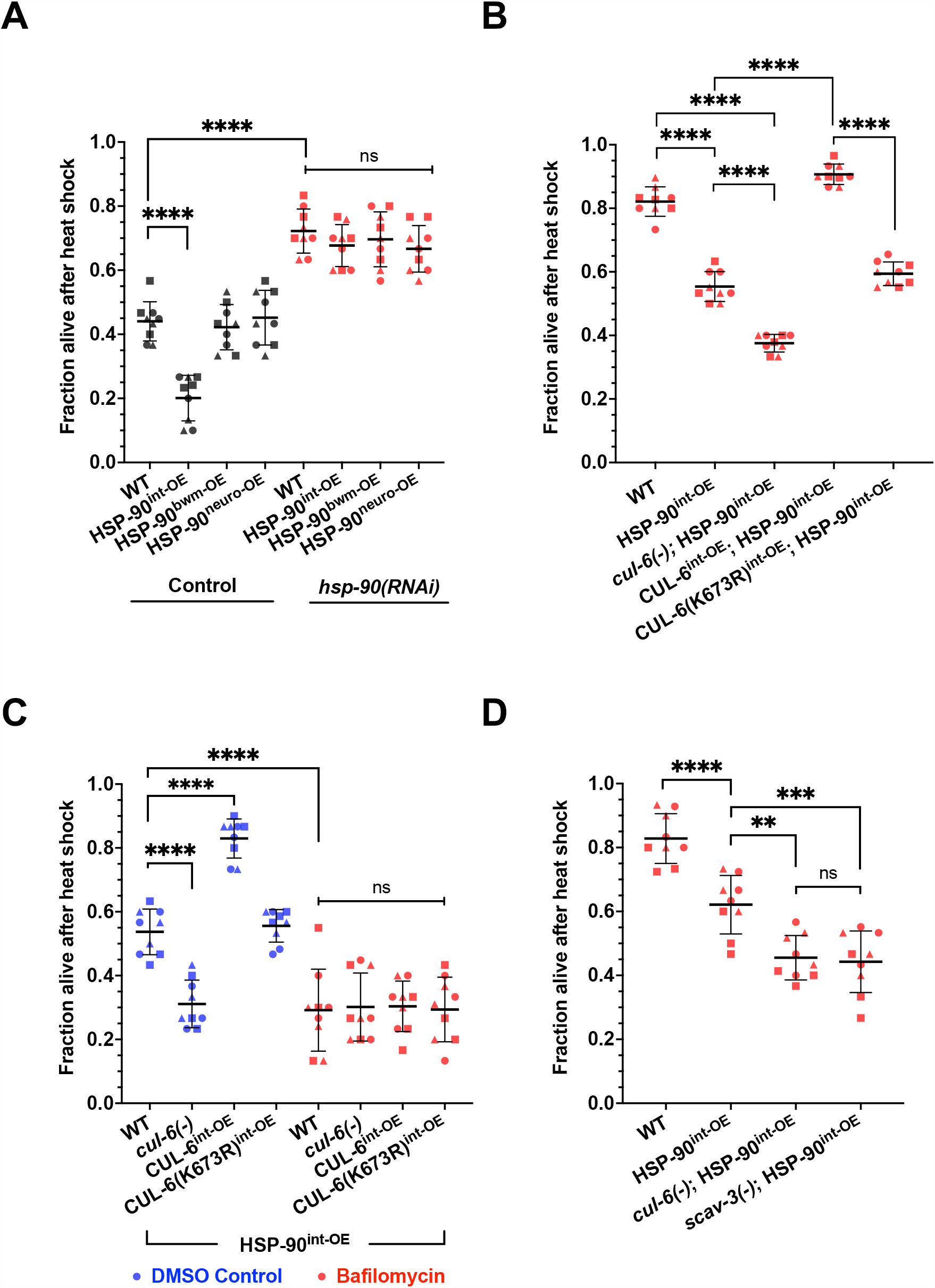
Overexpression of HSP-90 in the intestine decreases thermotolerance, which is reversed by CUL-6 expression and lysosome function. (A) Heat shock treatment of animals with HSP-90::RFP (hereafter HSP-90 for brevity) overexpression under the control of intestinal (superscript int-OE), body wall muscular (superscript bwm-OE), or neuronal (superscript neuro-OE) tissue-specific promoters fed OP50 expressing control (black) or *hsp-90* (red) dsRNA to induce RNAi. (B) Survival of wild-type, HSP goint-OE, and HSP-90^int-OE^ animals with *cul-6(-)* mutation or CUL-6^int-OE^ after reduced heat shock treatment (see methods). (C) Heat shock treatment of wild-type, *cul-6(-)*, or CUL-6^int-OE^ animals in an HSP-90^int-OE^ background treated with bafilomycin (red). DMSO-treated animals (blue) were tested as a vehicle control. (D) Survival of wild-type, HSP-90^int-OE^, and HSP-90^int-OE^ animals in a *cul-6(-)* or *scav-3(-)* mutant background after reduced heat shock treatment. For A-D, animals were tested in triplicate experiments with three plates per experiment and 30 animals per plate. The mean fraction of animals alive for the pooled replicates is indicated by the black bar with error bars as the SD. Each dot represents a plate, and different shapes represent the experimental replicates performed on different days. A one-way ANOVA with Tukey’s multiple comparisons test was used to calculate p-values. **p < 0.01, ***p < 0.001, ****p < 0.0001.

To confirm this intestinal-specific effect on thermotolerance was due to HSP-90 levels, we treated these tissue-specific expression strains with *hsp-90* RNAi and found that all of them had thermotolerance levels increased to the same level (Figure 3A). In fact, all of the overexpression strains had greatly increased thermotolerance after *hsp-90* RNAi, comparable with the survival rates after *hsp-90* RNAi in wild-type animals mentioned above (Figure 2A). Furthermore, intestinal HSP-90::RFP impairment of thermotolerance was not exacerbated by an *hsf-1* mutation, again suggesting that these effects are independent of HSF-1 (Figure S2A). To ensure that the effects on thermotolerance were specific to overexpression of HSP-90 in the intestine and not overexpression of the RFP tag, we tested thermotolerance of two other strains that specifically express cytoplasmic RFP in the intestine. We did not observe decreased thermotolerance in either of these strains (Figure S2B). Therefore, we conclude that overexpression of HSP-90 specifically in the intestine impairs thermotolerance.

### CUL-6 ubiquitin ligase and lysosomal activity promote thermotolerance when HSP-90 is overexpressed in the intestine

Next, we investigated the role of CUL-6 in promoting thermotolerance in the context of HSP-90 overexpression. Our previous studies found that loss of *cul-6* in a *pals-22* mutant background impaired thermotolerance, but the loss of *cul-6* in a wild-type background did not, suggesting that only when *cul-6* was upregulated does it have a role in thermotolerance.^7,15^ However, we reasoned that if a CUL-6 ubiquitin ligase targets HSP-90 protein, there would be increased potential for interaction between HSP-90 protein with even low levels of the CUL-6 ubiquitin ligase complexes (i.e. in a wild-type background) when HSP-90 is overexpressed. Therefore, we predicted we might see a decrease in thermotolerance upon loss of *cul-6* in wild-type animals when HSP-90 is overexpressed. Indeed, we found that when HSP-90 is overexpressed in the intestine, loss of *cul-6* caused a decrease in thermotolerance, to a level below that caused by HSP-90 overexpression alone (Figure 3B, S1B). Furthermore, we found that CUL-6 overexpression in the intestine increases thermotolerance in an HSP-90 overexpression background (Figure 3B, 3C). Importantly, overexpression of CUL-6(K73R) did not have an impact on thermotolerance when HSP-90 is also overexpressed, indicating that the effect of CUL-6 on thermotolerance is dependent on the neddylation site of CUL-6, as would be expected if thermotolerance were promoted by the activity of a CUL-6 ubiquitin ligase (Figure 3B, 3C).

The findings in Figure 3A-B suggest that thermotolerance is promoted by CUL-6-mediated degradation of HSP-90, which findings in Figure 1 suggested would occur in the lysosome. Indeed, we found that treatment with the lysosome inhibitor bafilomycin in an HSP-90 overexpression background reduced thermotolerance even further, suggesting that normally, lysosomal-mediated degradation of HSP-90 promotes thermotolerance (Figure 3C). Furthermore, treatment with bafilomycin suppressed the loss or overexpression of CUL-6, as would be expected if the lysosome were downstream of CUL-6 (Figure 3C). To further test the model that the lysosome degrades HSP-90 to promote thermotolerance, we crossed a *scav-3* mutation into the HSP-90 overexpression strain. Here, we found that a *scav-3* mutation causes reduced thermotolerance in an HSP-90 overexpression background (Figure 3D), while it does not affect thermotolerance in a wild-type background (Figure 1D). Furthermore, overexpression of SCAV-3::GFP,24 which has been shown to increase the function of lysosomes, leads to increased thermotolerance in an HSP-90 overexpression background (Figure S2C). Altogether, these results support the model that a CUL-6 ubiquitin ligase promotes the degradation of HSP-90 in the intestine by the lysosomes to promote thermotolerance.

### CUL-6 and lysosomal function regulate HSP-90 protein levels in the intestine

If CUL-6 ubiquitin ligases target HSP-90 for ubiquitylation and subsequent degradation by the lysosome, the levels of HSP-90 protein should be regulated by CUL-6 and lysosomal function. To investigate this possibility, we quantified HSP-90::RFP levels in strains with loss or overexpression of CUL-6. Because endogenous CUL-6 is expressed more highly in the anterior intestine compared to the rest of the intestine, we focused our quantification on the anterior half of the intestine. Here, we found that HSP-90::RFP levels varied inversely with CUL-6 activity. In particular, we saw significantly higher HSP-90::RFP levels in *cul-6* mutants compared to wild-type animals and significantly lower HSP-90::RFP levels in animals overexpressing wild-type CUL-6 specifically in the intestine, but not in those expressing neddylation-deficient CUL-6 (Figure 4A-4C).

**Figure 4.**
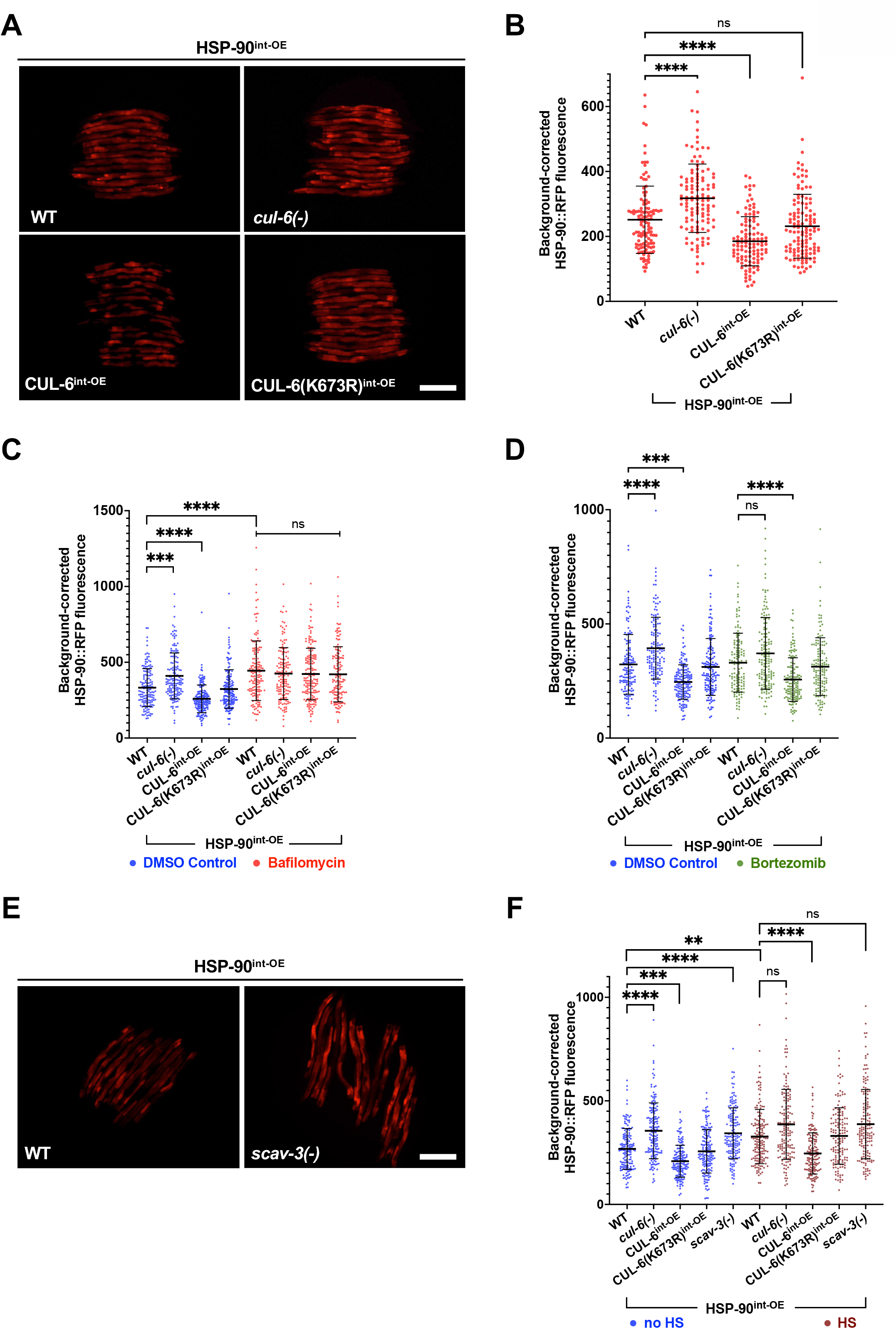
CUL-6 expression and lysosomal function reduce HSP-90 protein levels in the intestine. (A) Fluorescent images of L4 HSP-90^int-OE^ animals with mutation or overexpression of *cul-6*. Scale bar= 200 µm. (B) Quantification of RFP signal in the anterior intestine of strains shown in A. n = 113 - 125 worms quantified. (C) Quantification of RFP signal in the anterior intestine of L4 HSP-90^int-OE^ animals with mutation or overexpression of *cul-6* and treated with bafilomycin (red) or DMSO-(blue) as a vehicle control. n = 141 - 166 worms quantified. (D) Quantification of RFP signal in the anterior intestine of L4 HSP-90^int-OE^ animals with mutation or overexpression of *cul-* and treated with bortezomib (green) or DMSO (blue) as a vehicle control. n = 135-160 worms quantified. (E) Fluorescent images of wild-type or *scav-3(-)* mutant L4 animals expressing HSP goint-OE_Scale bar= 200 µm. (F) Quantification of RFP signal in the anterior intestine of HSP-90int oE animals with mutation or overexpression of *cul-6* in the absence of heat shock (blue) or 1 h after reduced heat shock treatment (maroon). n = 145 – 162 worms quantified. For B, C, D, and F, each dot on the graph represents one animal. The black bar indicates the mean fluorescence intensity with error bars as SD. A Kruskal-Wallis test with Dunn’s multiple comparisons test was used to calculate p-values; ****p <0.0001; *** p < 0.001; **p < 0.01.

To investigate the role of the lysosome in regulating HSP-90::RFP levels in the intestine, we treated HSP-90 overexpression animals with bafilomycin and then quantified HSP-90::RFP in the anterior intestine. Here, we saw higher levels of HSP-90 in animals treated with bafilomycin (Figure 4C). Furthermore, we found that bafilomycin treatment suppressed the effects of *cul-6* mutation or overexpression on HSP-90::RFP levels, consistent with the model that the lysosome acts downstream of a CUL-6 ubiquitin ligase to regulate HSP-90::RFP protein levels (Figure 4C). As a control for specificity, when we treated HSP-90 overexpression animals with the proteasome inhibitor bortezomib, we saw that *cul-6* mutation or overexpression still had an effect on HSP-90 levels (Figure 4D). We also investigated the role of the lysosome in regulating HSP-90::RFP levels by analyzing the effects of a *scav-3* mutation. Again, we saw that loss of lysosomal function in *scav-3* mutants led to increased levels of HSP-90 (Figure 4E and Figure 4F), comparable to the effect of loss of *cul-6* (Figure 4E). Furthermore, we found that *cul-6* and *scav-3* regulated levels of HSP-90 both before and after heat shock (Figure 4F). These results suggest that CUL-6 and lysosomes are able to degrade HSP-90 at both normal growth temperatures and during heat shock.

If HSP-90 protein were targeted for ubiquitylation and degradation, overexpression of ubiquitin might lower HSP-90 levels. Indeed, we found that overexpressing ubiquitin-GFP in intestinal cells led to lower HSP-90::RFP levels in the intestine (Figure S3). Ubiquitylation is the process of directly conjugating ubiquitin onto substrate proteins.^13^ Therefore as a control for non-specific effects of ubiquitin-GFP overexpression that are unrelated to ubiquitylation, we overexpressed a conjugation-deficient ubiquitin-GFP and here we saw no effect on HSP-90::RFP levels (Figure S3). Altogether, these findings support a model whereby wild-type *cul-6* promotes ubiquitylation of HSP-90, which leads to its degradation in the lysosome to promote thermotolerance.

### CUL-6 promotes HSP-90::RFP co-localization with lysosome-related organelles upon heat shock

To investigate if HSP-90 protein is targeted to the lysosome in a manner dependent on CUL-6, we imaged the subcellular localization of HSP-90::RFP before and after heat shock. Here, we saw that after heat shock, HSP-90::RFP localized to spherical structures in the intestine in a manner dependent on CUL-6; there were fewer spherical structures in *cul-6* mutants and more structures upon wild-type CUL-6 intestinal overexpression, but not upon neddylation-deficient CUL-6 intestinal overexpression (Figure 5). Based on fluorescence in the blue channel, these spherical HSP-90::RFP structures are localized to lysosome-related organelles, which are abundant organelles in the intestine with autofluorescence prominently in this wavelength.^34-37^ Lysosome-related organelles are multi-functional acidic compartments that express many canonical lysosomal markers, and have degradative potential.^38^ Thus, our results indicate that the CUL-6 ubiquitin ligase directs HSP-90::RFP to lysosomes and/or lysosome-related organelles for degradation upon heat shock in the intestine.

**Figure 5.**
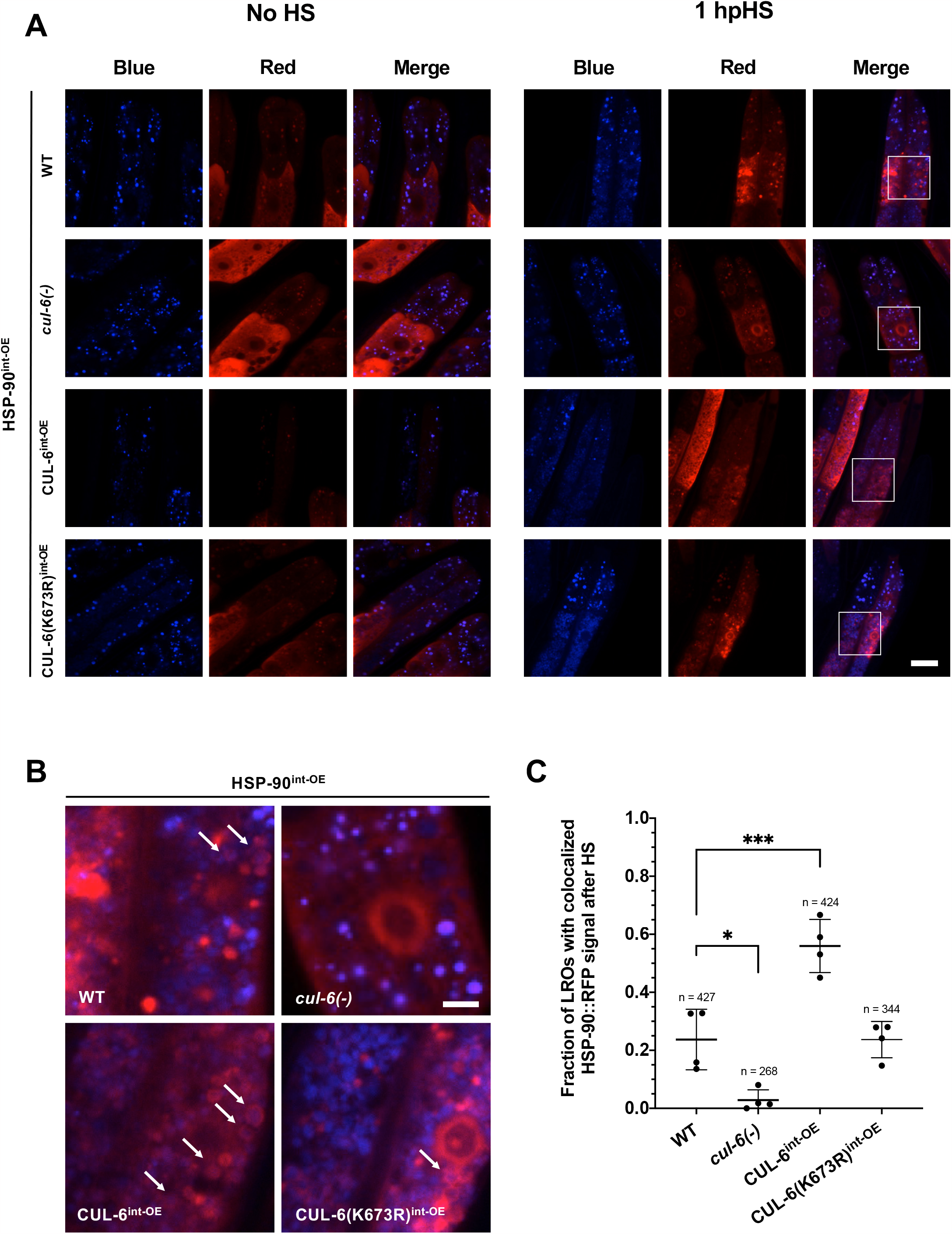
CUL-6 promotes HSP-90::RFP co-localization with lysosome-related organelles upon heat shock. (A) Representative confocal fluorescence images of the anterior intestine in L4 animals with varying levels of CUL-6 expression in strains in an HSP-90^int-OE^background both 1 h after reduced heat shock and in the absence of heat shock. Lysosome-related organelles (LROs) were imaged using blue channel autofluorescence signal. A region of interest showing colocalization events in the second ring of intestinal cells is highlighted in the merged image 1 h after reduced heat shock. Scale bar= 20 µm. (B) Enlarged regions of interest identified in panel A showing colocalization of HSP-90 to LROs. Arrows depict ring-like aggregation pattern of HSP-90::RFP upon heat shock around LROs. Scale bar = 5 µm. (C) Quantification of the fraction of LROs with colocalized HSP-90 events after reduced heat shock treatment in the second ring of intestinal cells. n = 4 worms quantified, representing 268 – 427 LROs per condition; results representative of two independent experiments. A one-way ANOVA with Tukey’s multiple comparisons test was used to calculate p values. *p < 0.05, ***p < 0.001.

## Discussion

Previously, we identified the IPR-regulated CUL-6 ubiquitin ligase as a novel proteostasis pathway in *C. elegans* that is upregulated by intracellular infection of the intestine and by proteotoxic stress including chronic heat stress. In this study, we identified HSP-90 as a target of the CUL-6 ubiquitin ligase in defending the host from proteotoxic stress. In particular, we show that CUL-6 promotes degradation of HSP-90 protein by lysosomes and/or lysosome-related organelles in the intestine (Figure 6). Several other studies have shown that HSP-90 can bind to and negatively regulate HSF-1, a master regulator transcription factor of the heat shock response.^22,23,39^ Thus, we considered whether CUL-6-mediated degradation of HSP-90 may increase thermotolerance through activation of HSF-1. However, we found that CUL-6-mediated degradation of HSP-90 appears to promote thermotolerance independently of HSF-1, a result consistent with our initial characterization of the IPR being independent of the canonical heat shock response mediated by HSF-1.^7^

**Figure 6.**
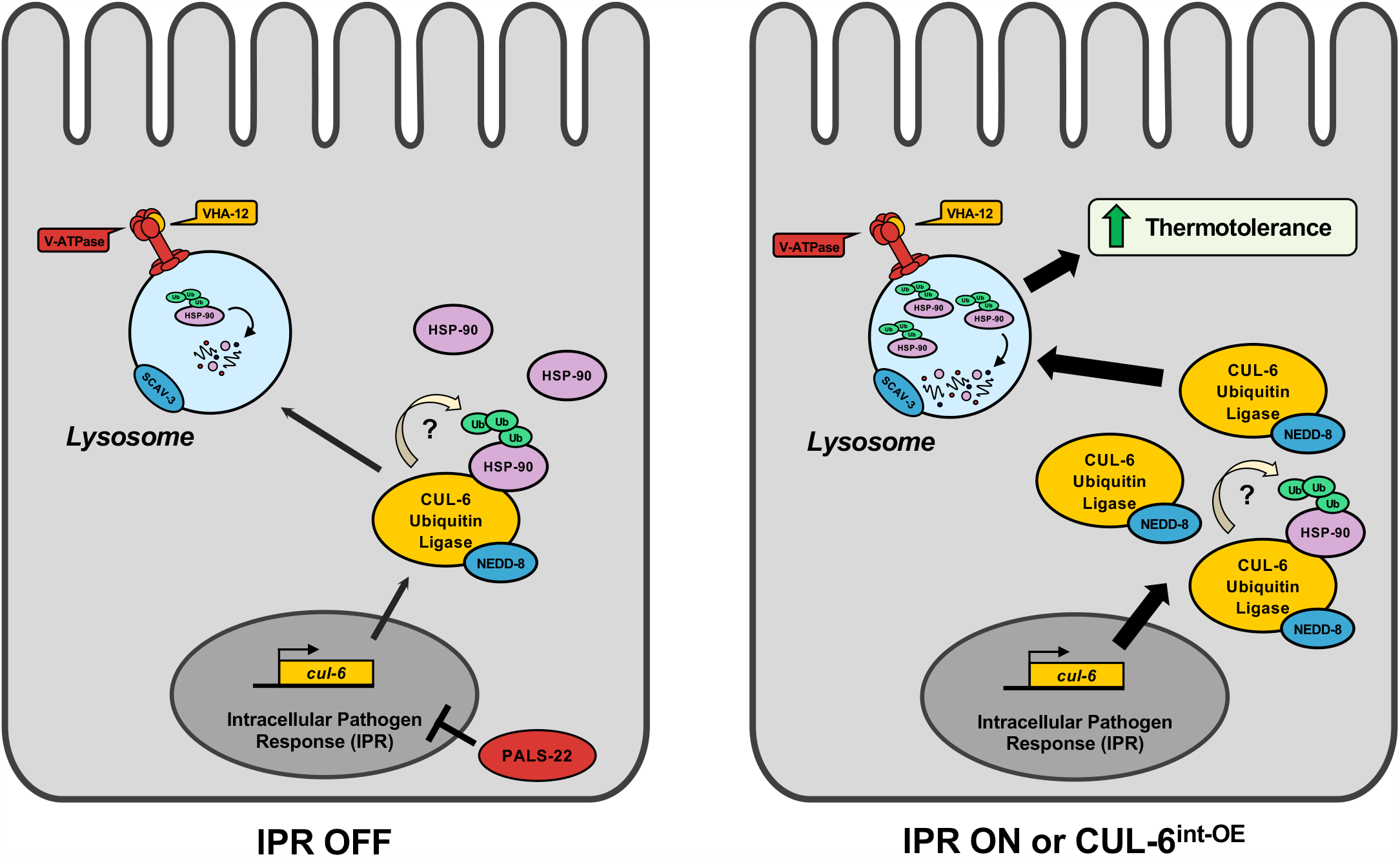
A model for CUL-6-mediated degradation of HSP-90. The cullin CUL-6 is upregulated as part of the IPR, a novel stress response pathway. HSP-90 is a substrate of the CUL-6 ubiquitin ligase complex and is degraded in the lysosome. Degradation of HSP-90 improves survival after heat shock stress, or thermotolerance. When the IPR is activated, upregulated expression of CUL-6 leads to increased trafficking of HSP-90 to the lysosome, resulting in higher thermotolerance relative to an inactive IPR state.

Identifying the CUL-6-mediated reduction of HSP-90 protein levels provides insights into the mechanisms by which the IPR promotes thermotolerance. Degradation of a heat shock protein to promote thermotolerance (independently of HSF-1) may seem counterintuitive because heat shock proteins as a protein family canonically promote thermotolerance, but it is important to note that HSP-90 is distinct from other heat shock proteins. HSP-90, together with its many co chaperones, is a central player in protein folding in the absence of heat shock. Unlike other heat shock proteins, HSP-90 is highly expressed in the absence of heat shock, and in fact the HSP-90 protein itself is thought to comprise 1-2% of the entire proteome under unstressed conditions^19^. Due to the high basal level of HSP-90 expression, even the modest ∼25% increase or decrease of HSP-90 levels caused by loss or overexpression of CUL-6 (Figure 4) may actually be a substantial change in the absolute amount of this highly abundant protein.

HSP-90 plays a broad role in facilitating folding and maturation of the proteome and in fact is estimated to be required for the activity of up to 20% of all proteins and 60% of all kinases^40,41^ .Among its varied roles, HSP-90 promotes the active conformation of oncogenic kinases, aids assembly of the multi-protein kinetochore complex, and promotes ligand binding to steroid hormone receptors^19^. All of these HSP-90 clients (proteins whose folding is facilitated by HSP-90) are important for cellular growth in the absence of heat shock. Therefore, one hypothesis to explain our findings is that degradation of HSP-90 in the *C. elegans* intestine leads to increased thermotolerance because it prevents maturation and/or degrades clients of HSP-90 that promote growth in the intestine. In this way, a CUL-6 ubiquitin ligase could promote the removal of multiple proteins at once after heat shock, and provide a ‘factory reset’ away from a growth state and toward a reparative state. However, altered levels of other HSP-90-associated proteins, as well as other distinct mechanisms, could explain our results showing that CUL-6-mediated degradation of HSP-90 promotes thermotolerance.

A simple model for our results is that degradation of HSP-90 in the intestine alone can promote organismal thermotolerance, but other tissues may also be involved. Studies of systemic signaling in stress responses have shown that RNAi knock-down of *hsp-90* mRNA in either the intestine or in neurons triggers upregulation of *hsp-70* mRNA expression in muscle cells to promote thermotolerance, a phenomenon named ‘transcellular chaperone signaling’.^20,42,43^ These effects were recently found to be HSF-1-independent, as are the effects we see here with CUL-6. Therefore, some of the thermotolerance benefits from CUL-6-mediated loss of HSP-90 protein may be due to cell non-autonomous signaling to muscle. In fact, the IPR may provide a physiologically relevant stimulus to explain the activation of transcellular chaperone signaling, which has previously been studied through RNAi knock-down. Perhaps this form of systemic signaling is normally triggered by natural intracellular infection of the intestine.

While our model posits that a CUL-6 ubiquitin ligase ubiquitylates HSP-90 to send it to the lysosome for degradation, we have not directly shown this ubiquitylation event. Demonstrating this event will likely be challenging because of the modest relative change in HSP-90 levels (Figure 4). In addition, there are at least 7 subunits to the CUL-6 ubiquitin ligase complex(es),^15^ making biochemical reconstitution of this ligase together with candidate substrates difficult. Furthermore, HSP-90 has 42 lysines that are predicted ubiquitylation sites (as assessed by prediction software on this site: https://www.biocuckoo.org/), making it challenging to determine which sites are important for ubiquitylation. With these ideas in mind, a direct investigation of HSP-90 ubiquitylation by a CUL-6 ubiquitin ligase will be part of future studies.

While HSP-90 functions to fold its client proteins, it also aids in their degradation.^44-47^ To our knowledge, this study is the first to show degradation of HSP-90 itself, by the lysosome. Numerous drugs inhibit HSP-90, such as those developed to treat cancer that lead to degradation of oncogenic kinase clients of HSP-90.^48^ Thus, if the pathway we have identified for degradation of HSP-90 itself in *C. elegans* is conserved in humans, our findings may have relevance for efficacy of those drugs. Several studies in *C. elegans* have now shown that reduction of protein quality control factors can paradoxically promote organismal survival. In addition to the transcellular chaperone signaling described above, repression of *hsp-70* mRNA by microRNA after heat shock has been shown to aid in heat shock recovery.^49^ Another example is a recent study showing that inhibition of HSF-1 causes decreased protein aggregation and better functioning specifically of the pharynx.^50^ Interestingly, this protective benefit in the pharynx is associated with upregulation of IPR genes and relies on the same lysosomal factors we found were important for the thermotolerance benefit of CUL-6-mediated degradation of HSP-90 in our study. In the future, it will be exciting to determine the connection among all these findings, and to uncover the downstream mechanism(s) by which degradation of the central proteostasis factor HSP-90 promotes organismal survival after heat shock.

## Materials and Methods

### C. elegans maintenance and strain generation

Worms were maintained using standard methods at 20 °C on Nematode Growth Media (NGM) agar plates top-plated with streptomycin-resistant *Escherichia coli* OP50-1 unless stated otherwise.^51,52^ Worm strains used in this study are listed in Table S1.

### Synchronization of C. elegans

To obtain synchronized populations of L1 *C. elegans* for fluorescent imaging experiments described below, gravid adult animals were collected from NGM+OP50-1 plates in M9 buffer into a 15 mL conical tube. The tubes were centrifuged, and the supernatant was removed, leaving animals in ∼2 mL of M9. 800 µL of 5.65-6% sodium hypochlorite solution and 200 µL of 2M NaOH were added to the tube, and the contents were vigorously shaken for approximately 1 minute and 40 seconds. Embryos released after bleaching were resuspended in 15 mL of M9, centrifuged, and the supernatant was discarded. The embryos were washed with M9 in this manner a total of 5 times, resuspended in a final volume of 5 mL of M9, and then placed in a 20 °C incubator under continuous rotation for 16-24 hours until L1s hatched.

### Thermotolerance Assays

Heat shock treatment: Gravid adults were picked to 6-cm NGM+OP50-1 plates and grown at 20 °C. 30 F1 progeny from these adults were picked at the L4 life stage to fresh NGM+OP50-1 plates and subjected to a heat shock of 37.5 °C in a dry programmable incubator for 2 hours and 15 minutes with an initial gradual ramp-up to 37.5 °C over a 15 minute period (2 hours stable at 37.5 °C). Immediately following the completion of the 2-hour and 15 minute heat shock program, the plates were removed from the incubator. The animals were allowed to recover for 30 minutes at room temperature by placing the plates in a single layer on a benchtop, then incubated at 20 °C for 24 hours. Animals were then scored for survival in a blinded manner; worms not responding to touch with a worm pick, defined by a single prod to the body, were scored as dead. 3 replicate plates were scored for each condition per experiment, and the experiment was performed 3 independent times, except as noted for the DMSO and bortezomib treatment experiment performed in Figure 1A. A summary of the heat shock treatment and experiments where it was applied in this study is provided in Figure S1A.

Reduced heat shock treatment: A modified version of the assay with a shorter duration was used to accommodate the increased heat shock susceptibility of the HSP-90::RFP^int-OE^ strain. Following the standard protocol described above, most HSP-90::RFP^int-OE^ animals died. Therefore, to improve survival and allow for the analysis of HSP-90 intestinal overexpression on raising or lowering thermotolerance phenotypes in different contexts, the assay time was shortened to 2 hours of 37.5 °C heat shock with a 15 min gradual ramp-up (1 hour 45 minutes stable at 37.5 °C). Immediately following the completion of the 2-hour heat shock program, the plates were removed from the incubator. Recovery, scoring, and experimental replicates were performed as described above. A summary of the reduced heat shock treatment and experiments where it was applied in this study is provided in Figure S1B.

### RNAi experiments

RNAi was performed by the feeding method.^53^ Overnight cultures of OP50 strain (R)OP50, modified to enable RNAi^54^ were plated on 6-cm RNAi plates (NGM plates supplemented with 5mM β-D-1-thiogalactopyranoside [IPTG], 1 mM carbenicillin), and incubated at 20 °C for 3 days. Gravid adults were transferred to these plates for growth. The F1 progeny at the L4 stage were transferred to new, matching 6-cm RNAi plates before being tested for thermotolerance as previously described. OP50 RNAi strains were generated by extracting the desired RNAi plasmid vector from HT115 *E. coli* in the existing Ahringer and Vidal RNAi libraries. The plasmids were then transformed into competent (R)OP50, and transformants were selected after -24 hours of growth based on carbenicillin resistance. Overnight cultures of transformations were then mini prepped, and the L4440 plasmid vector was sequenced with the T7 forward primer to confirm that the insert matching the desired gene for knockdown studies was present in the vector.

### Bortezomib Treatment

The proteasome was inhibited with bortezomib (Selleckchem Chemicals LLC, Houston, TX) as previously described.^11^ For thermotolerance assays, gravid adults were plated onto 10-cm NGM+OP50-1 plates and incubated at 20 °C for 72 hours. Before performing heat shock treatment, a 10 mM stock solution of bortezomib in DMSO was top-plated to 6-cm NGM+OP50-1 plates to reach a final concentration of 2 µM, 10 µM, or 20 µM per plate, in duplicate per condition. The same volume of DMSO was added to the control plates. The plates were dried, and 30 L4 F1 animals were transferred onto each treatment or control plate. The plates were then subjected to the standard heat shock treatment, recovery, and scoring regimen described above.

To quantify HSP-90::RFP fluorescence, synchronized L1s were plated on 10 cm NGM+OP50-1 plates and grown for 44 h at 20 °C. A 10 mM stock solution of bortezomib in DMSO was top-plated to reach a final concentration of 20 µM per plate, and the same volume of DMSO was added to control plates. Plates were dried, and worms were incubated for 4 h at 20 °C. The animals were washed off the treatment plates in M9+tween-20, pelleted, anesthetized with a final concentration of 10mM levamisole, and transferred into a 96-well plate with fresh 10 mM levamisole in M9+tween-20. Imaging was performed using the lmageXpress Nano using a 4x objective (Molecular Devices, LLC) and analyzed using the FIJI program. Quantification of fluorescent signal was restricted to the anterior intestine, defined for the purposes of this assay as the beginning of the intestine to the midpoint of the vulva region, since CUL-6 is primarily expressed in the anterior intestine and we would expect that mutation of endogenous CUL-6 would have the greatest effect on HSP-90 levels in this region. The background signal was collected from three adjacent regions and the mean subtracted from the signal measured in the anterior intestines of animals.

### Bafilomycin Treatment

The lysosome was inhibited with bafilomycion A1 (AdipoGen Life Sciences, Liestal, Switzerland). For thermotolerance assays, a 25 mM stock solution of bafilomycin in DMSO was added to 6-cm NGM+OP50-1 plates to reach a final concentration of 1 µM or 3 µM per plate in triplicate per condition. The same volume of DMSO was added to the control plates. Plates were dried, gravid adults added, and the plates were then incubated for 72 hours at 20 °C. 30 L4 F1 animals from each plate were then transferred onto freshly prepared of 3 µM and dried plates with matching treatments. The plates were then subjected to the standard heat shock treatment, recovery, and scoring regimen described above.

To quantify HSP-90::RFP fluorescence, a 25 mM stock solution of bafilomycin in DMSO was top plated to a 10-cm NGM+OP50-1 plate to reach a final concentration of 3 µM per plate. The same volume of DMSO was added to the control plates. Once the plates were dry, synchronized L1s were added, and the plates were incubated for 48 hours at 20 °C. The animals were washed off the treatment plates in M9+tween-20, pelleted, anesthetized with a final concentration of 10mM levamisole, and transferred into a 96-well plate with fresh 10 mM levamisole in M9+tween-20. Imaging was performed using the lmageXpress Nano using a 4x objective (Molecular Devices, LLC) and analyzed using the FIJI program. Quantification was restricted to the anterior intestine and background-corrected as described above.

### Microscopy

For HSP-90::RFP-expressing animals shown in Figures 4A, 4E, and S2A, L4 animals were picked from a mixed population grown on 6-cm NGM+OP50-1 plates incubated at 20 °C, anesthetized with 10 mM levamisole in M9 buffer, and mounted on 2% agarose pads for imaging on a Zeiss Axioimager M1 compound microscope with a 10X objective. For S2A, whole animal fluorescent signal was quantified using the FIJI program, corrected by the mean background sampled from three adjacent background regions.

For images of HSP-90::RFP in the absence of and 1 hour after heat shock (Figure 5), animals were synchronized by bleaching as described above, then cultured on 10 cm NGM+OP50-1 plates for 44 hours at 20 °C. While control (no heat shock) animals remained at 20 °C, animals assigned to heat shock treatment were subjected to a reduced heat shock exposure (2 hours with a 15-minute ramp up to 37.5 °C, 1 hour and 45 minutes stable at 37.5 °C), and allowed to recover for the standard 30 minutes at room temperature. Plates were then incubated at 20 °C for 1 hour before all conditions were anesthetized with 10 mM levamisole in M9 buffer and mounted on 2% agarose pads for imaging with a Zeiss LSM700 confocal microscope and Zen2010 software using a 63X objective. For quantification, the fraction of LROs in the second ring of intestinal cells as labeled by autofluorescence in the blue channel with positive HSP-90::RFP colocalization over total number of LROs in the region of interest was recorded for a total of four animals per condition.

### Statistics

All statistical analysis was performed with Prism 9 software (GraphPad). The D’Agostino & Pearson omnibus normality test was used to assess the data distribution for all experiments. Standard parametric statistical tests were performed for normally distributed data, and nonparametric tests were performed for non-normally distributed data. The corresponding figure legends describe the statistical test used for each experiment, the number of data points analyzed, replicate information, and other relevant details.

## Acknowledgements

We thank Michalis Barkoulas, Lakshmi Batachari, Aundrea Koger, Katie Li, Johan Panek, Deevya Raman and Nicole Wernet for helpful comments on the manuscript. Some strains used in this study were provided by the Caenorhabditis Genetics Center (CGC), which is funded by NIH Office of Research Infrastructure Programs (P40 OD010440). This work was supported by NIH under R01 AG052622, GM114139, Al176639 and by NSF 2301657 to E.R.T., by NIGMS/NIH award K12GM068524 to S.S.G. and by NIH R01 AG082970 to P.v.O.-H.

## Supplemental Figures

**Figure S1.**
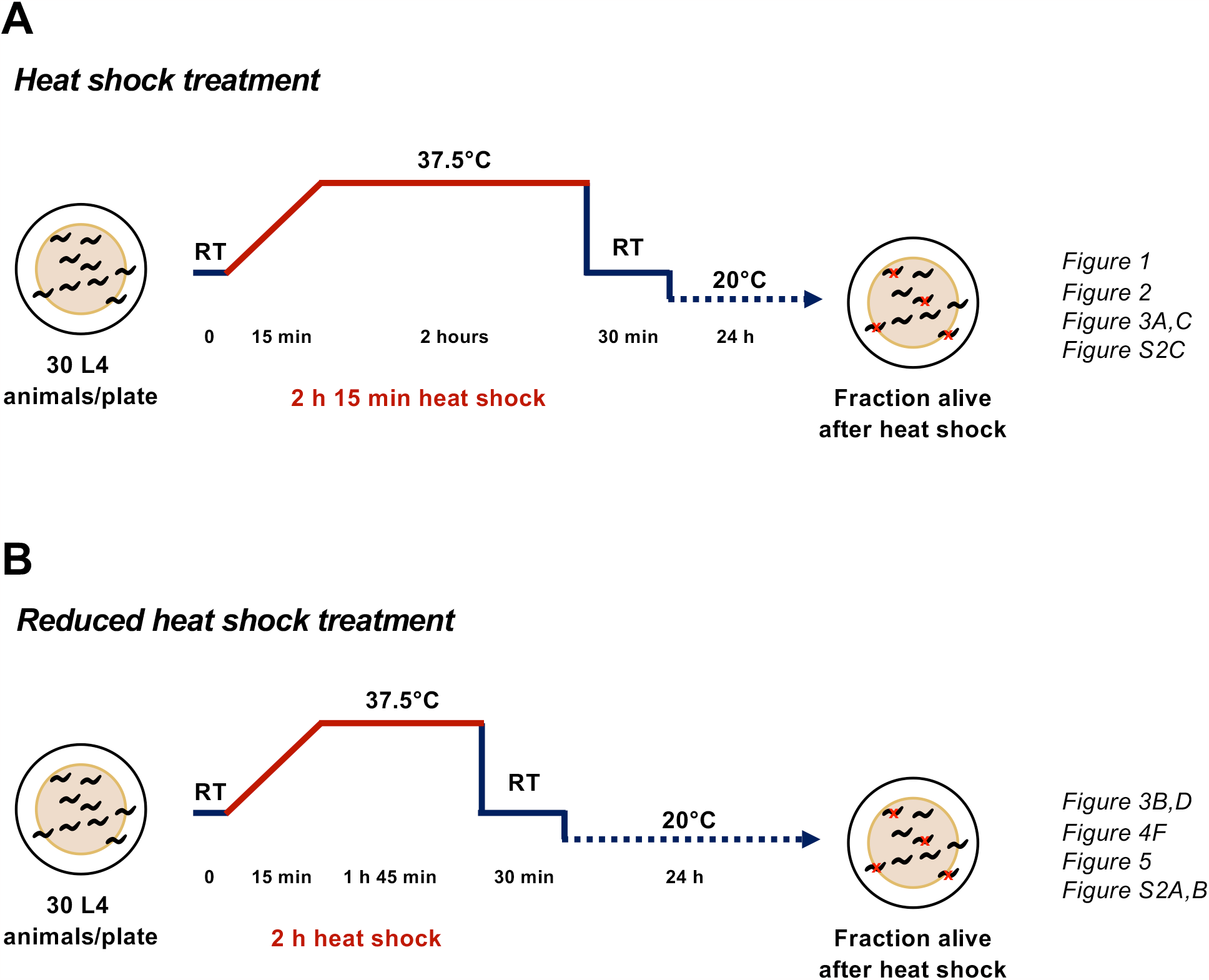
Heat shock treatments to assess thermotolerance phenotypes. (A) Diagram of the workflow for the 2 h 15 min “heat shock treatment” and experiments where it was applied (also see methods). (B) Diagram of the workflow for the 2 h “reduced heat shock treatment” and experiments where it was applied (also see methods). HSP-90^int-OE^ strains survive poorly following the longer heat shock treatment shown in A (example: Figure 3A, control condition). When warranted, strains with HSP-90^int-OE^ were instead subjected to the less stressful reduced heat shock treatment to better assess CUL-6 and lysosome-mediated phenotypes.

**Figure S2.**
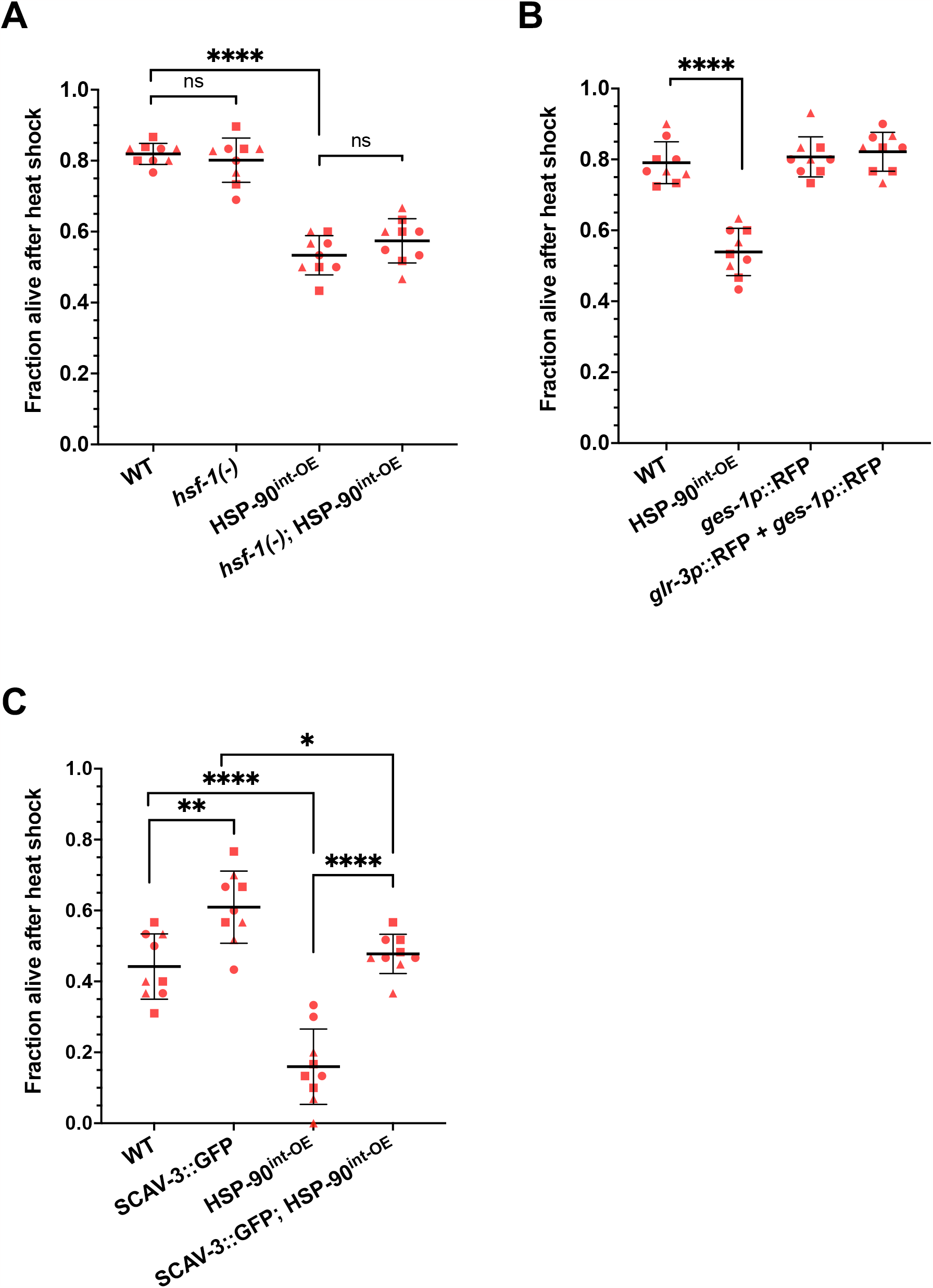
CUL-6 lowers HSP-90::RFP levels independent of HSF-1, and overexpression of SCAV-3::GFP promotes thermotolerance. (A) Survival of wild-type, *hsf-1(-)* mutants without or with HSP-90^int-OE^, and HSP-90^int-OE^ animals after reduced heat shock treatment. (B) Survival of wild-type, HSP-90^int-OE^ animals, and two additional strains that overexpress RFP in the cytosol of the intestine after reduced heat shock treatment. (C) Survival of wild-type and HSP-90^int-OE^ animals with or without SCAV-3::GFP overexpression after heat shock treatment. For A-C, animals were tested in triplicate experiments with three plates per experiment and 30 animals per plate. The mean fraction of animals alive for the pooled replicates is indicated by the black bar with error bars as the SD. Each dot represents a plate, and different shapes represent the experimental replicates performed on different days. A one-way ANOVA with Tukey’s multiple comparisons test was used to calculate p-values.; ****p<0.0001; **p < 0.01; *p < 0.05; ns indicates no significant difference.

**Figure S3.**
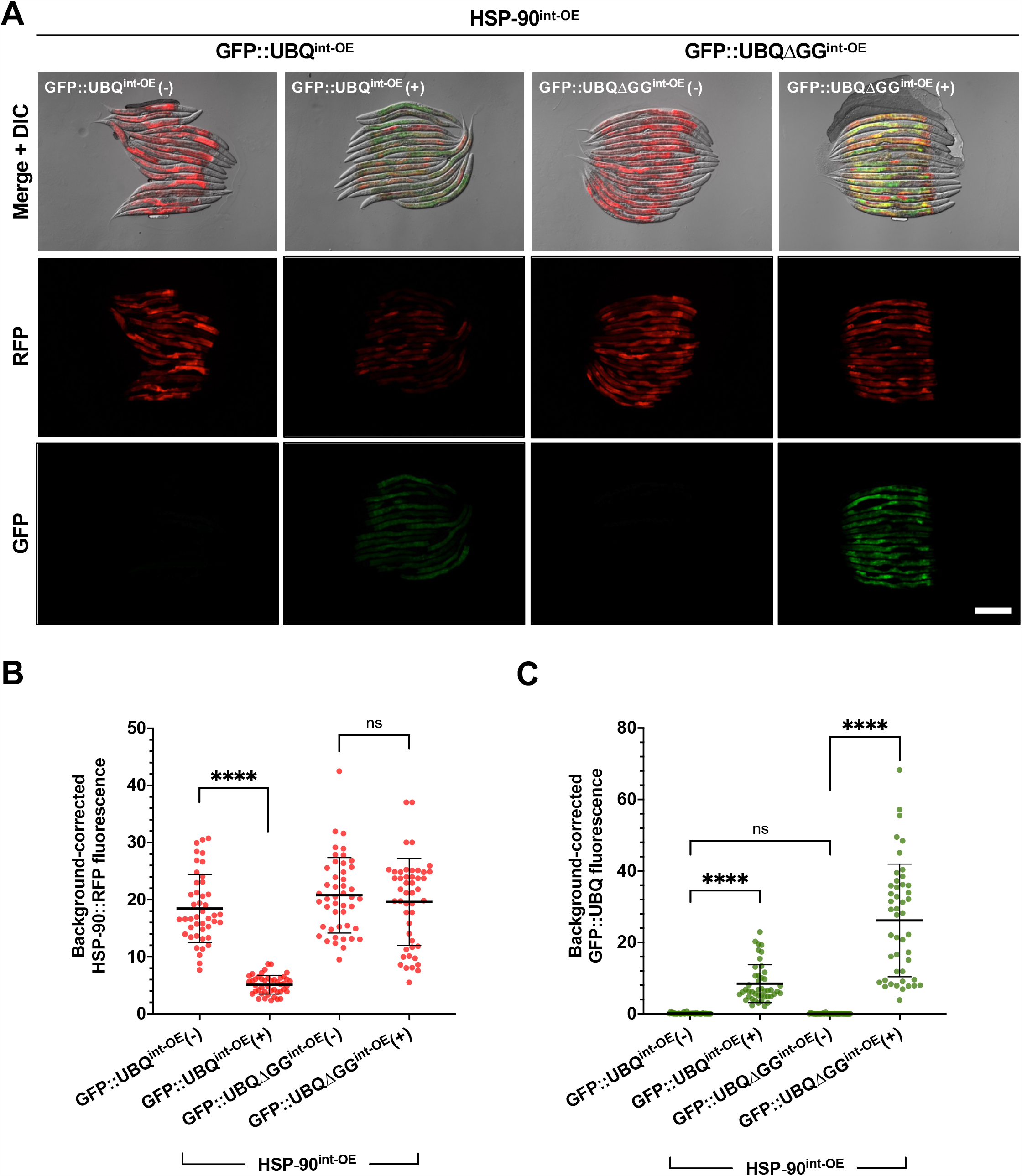
Ubiquitin overexpression in the intestine reduces H5P-90::RFP expression. (A) Representative fluorescent images of L4 animals carrying an extrachromosomal array that intestinally overexpresses either functional GFP::UBQ or a conjugation-deficient version, GFP::UBQΔGG, in an HSP-90^int-OE^ background. Sibling transgenic animals carrying both the GFP::UBQ/GFP::UBQΔGG array and HSP-90^int-OE^ or just HSP-90^int-OE^ alone are shown. Scale bar = 200 µm. (B) Quantification of the RFP fluorescent signal for each condition. (C) Quantification of the GFP fluorescent signal for the same animals measured in B. For B and C, each dot represents one animal measured, and the data shown are the results of three experimental replicates. n = 44 - 47 worms quantified. The black bar indicates the mean fluorescence intensity with error bars as SD. For each, a Kruskal-Wallis test was used to calculate p-values; ****p <0.0001.

## Supplemental Tables

**Table S1.**
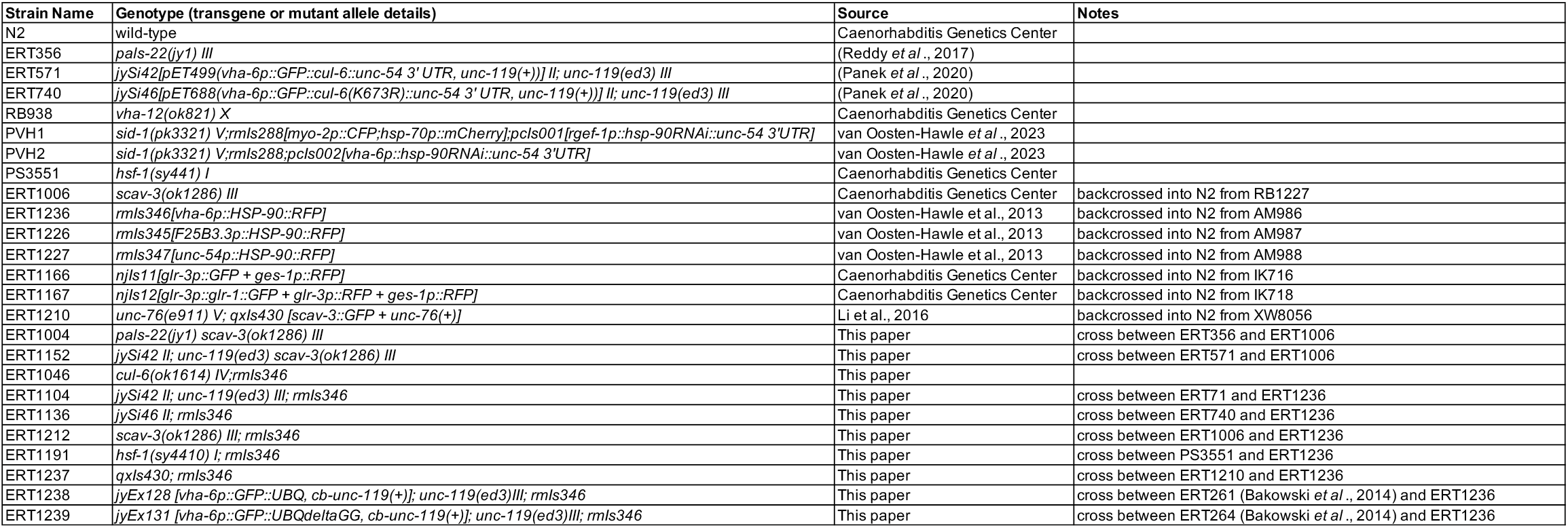
*C. elegans* strain list.

